# Single-cell sequencing of mouse thymocytes reveals mutational landscape shaped by replication errors, mismatch repair and H3K36me3

**DOI:** 10.1101/816207

**Authors:** Elli-Mari Aska, Denis Dermadi Bebek, Liisa Kauppi

## Abstract

**Background:** DNA mismatch repair (MMR) safeguards genome stability by correcting errors made during DNA replication. *In vitro* evidence indicates that the MMR machinery is recruited to chromatin via H3K36me3, a histone mark enriched in 3’ exons of genes and associated with transcriptional activity. To dissect how replication errors, abundance of H3K36me3 and MMR together shape the mutational landscape in normal mammalian cells, we applied single-cell exome sequencing to thymic T cells isolated from MMR-proficient (*Mlh1*^+/+^) and MMR-deficient (*Mlh1*^−/−^) mice.

**Results:** Using single-cell exome sequencing we identified short deletions as sensitive and quantifiable reporters of MMR-dependent mutations. We found H3K36me3-enriched *Huwe1* and *Mcm7* genes to be mutational hotspots exclusive to *Mlh1*^−/−^ T cells. In *Mlh1*^+/+^ cells, exons of H3K36me3-enriched genes had a lower mutation frequency compared to H3K36me3-depleted genes. Moreover, within transcriptionally active genes, 3’ exons, often H3K36me3-enriched, rather than 5’ exons had significantly fewer MMR-dependent mutations, indicating that MMR operates more efficiently within 3’ exons in *Mlh1*^+/+^ T cells.

**Conclusions:** Our results provide evidence that H3K36me3 confers preferential MMR-mediated protection from transcription-associated deleterious replication errors. This offers an attractive concept of thrifty MMR targeting, where genes critical for the development of given cell type are preferentially shielded from *de novo* mutations by H3K36me3-guided MMR.

## BACKGROUND

Maintaining genomic integrity during DNA replication is crucial for cellular homeostasis, especially in protein-coding regions. Occasionally, DNA replication errors occur, of which most, but not all, are corrected by the intrinsic proofreading activity of DNA polymerases (1). DNA mismatch repair (MMR) corrects base-base mismatches and small insertion-deletion (indel) loops that have escaped proofreading, and thereby protects the genome from replication induced permanent mutations (2). MMR initiates when the MSH2/MSH6 (MutSα) or MSH2/MSH3 (MutSβ) complex recognizes and binds DNA lesions, a step followed by recruitment of the MLH1/PMS2 (MutLα) complex that triggers the excision and repair of the mismatch (3, 4).

MSH6 of MutSα can bind to trimethylated histone H3 lysine 36 (H3K36me3) and recruit the MMR machinery to chromatin (5). H3K36me3 is found in exonic regions and enriched at the 3’ ends of transcribed genes (6), but also in constitutive and facultative heterochromatin (7). Genome-wide mutational analyses of MMR-deficient cell lines and tumors have shown that presence of H3K36me3 reduces local mutation rate (8, 9). Moreover, MMR operates more efficiently in H3K36me3-enriched exons compared to introns (10), and in actively transcribed genes compared to silent genes (11).

MMR deficiency has been extensively modeled in *Mlh1*^−/−^ mice, which display high microsatellite instability (MSI) and increased mortality due to lymphomas and/or gastrointestinal tumors (12–15). MSI occurs due to the propensity of microsatellites (short tandem repeat sequences) to undergo strand slippage during DNA replication, which in MMR-deficient cells leads to deletion or insertion mutations within repeats. Recently, analysis of genome-wide mutations in *Mlh1*^−/−^ T cell lymphomas revealed several putative drivers of tumorigenesis (16).

To delineate how the mutational landscape in normal mammalian cells is shaped, on one hand, by replication errors, and on the other hand, by H3K36me3-mediated MMR correction, we performed single-cell whole exome sequencing (scWES) on T cells isolated from MMR-proficient (*Mlh1*^+/+^) and MMR-deficient (*Mlh1*^−/−^) mice. Comparison of mutation distribution and frequency between MMR-proficient and -deficient mice revealed *Huwe1* and *Mcm7* genes as mutational hotspots exclusive to *Mlh1*^−/−^ cells, implying that these regions present an inherent challenge to faithful DNA replication in T cells. Both hotspots are located in H3K36me3-enriched regions and expressed during T cell development. Analysis of MMR-dependent mutations indicate that H3K36me3-enriched 3’ exons are more protected against transcription-associated replication errors.

## RESULTS

### Deletions report on MMR-dependent mutations in single-cell exome sequencing

We isolated naïve T cells from thymi of *Mlh1*^+/+^ and *Mlh1*^−/−^ mice, followed by single-cell capture and whole genome amplification on the Fluidigm C1 system, and then, by whole exome enrichment and sequencing (Fig. 1). Previous studies have utilized single-cell DNA sequencing to study clonality and mutation profiles of human cancers and normal cells (17–20). To check whether T cells were drawn from a similar cell population in both genotypes, we analyzed the proportions of distinct developmental thymic T cell populations (double-negative (DN), double-positive (DB), TCR αβ single-positive (CD4 or CD8), TCR γδ) (21) by FACS. Cell frequencies of different thymic T cell populations between *Mlh1*^−/−^ and *Mlh1*^+/+^ mice were similar to each other (**Fig. S1**), indicating no defect in normal T cell developmental progression in *Mlh1*^−/−^ mice, and that T cells analyzed by scWES from *Mlh1*^+/+^ and *Mlh1*^−/−^ mice are drawn from similar thymic T cell populations. In both genotypes, the vast majority of cells were CD4+CD8+ double positive T cells (67% for *Mlh1*^+/+^ and 65% for *Mlh1*^−/−^ mice, respectively, **Fig. S1**).

**Figure 1.**
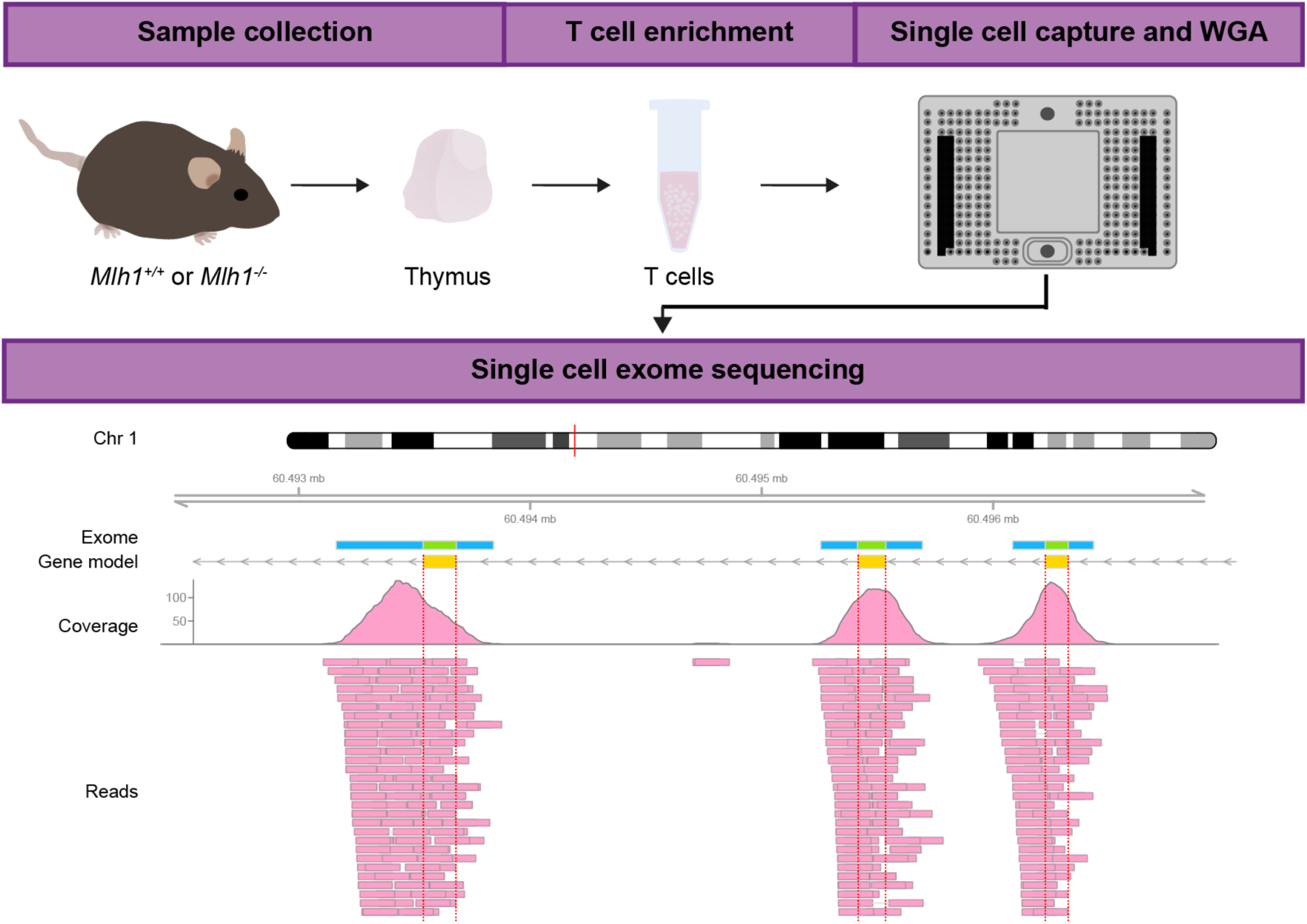
Whole exome sequencing of single T cells: Experimental overview. Thymi of *Mlh1*^−/−^ and *Mlh1*^+/+^ mice were dissected and used for enrichment of naïve T cells, followed by single-cell capture, cell lysis, and whole genome amplification in a Fluidigm C1. Amplified genomes were used for whole exome sequencing (WES) and sequencing reads were analyzed for genetic variants. Shown is a read pileup and coverage of sample WT1-C26 in a ~5-kb long region on chromosome 1 that contains three exons of *Raph1*. In addition to exons (green bar in exome panel), WES also partially covers non-coding regions adjacent to exons (blue bar in exome panel), enabling the comparison of mutation frequency between exonic and non-coding regions.

We sequenced 56 single-cell exomes in total, from 28 *Mlh1*^−/−^ and 28 *Mlh1*^+/+^ T cells, to an average depth of 32X and coverage of 66% at depth ≥1X (**Fig. S2A-B**). After excluding samples with low (< 50%) coverage, 44 exomes (22 *Mlh1*^+/+^ and 22 *Mlh1*^−/−^ exomes) were further analyzed for genetic variants. All detected variants with annotations are listed in Additional File 1.

Overall, *Mlh1*^−/−^ T cells had an increased percentage (O.R = 1.56, 95% CI = 1.44-1.69, p < 2.2×10^−16^) and frequencies (p = 5.487×10^−6^, Fig. 2A-B, Table S1) of indels when compared to *Mlh1*^+/+^ T cells. Even though MMR-deficiency increases also base substitutions (22), in our data set SNV frequencies between *Mlh1*^−/−^ and *Mlh1*^+/+^ did not differ significantly (p = 0.127, Fig. 2B, Table S1). Analyzing insertions and deletions separately revealed that *Mlh1*^−/−^ T cells had significantly higher deletion (p = 8.175×10^−12^), but not insertion frequencies (p = 0.1801) than *Mlh1*^+/+^ T cells (Fig. 2C, Table S1). Taken together, deletions behaved in a genotype-dependent manner, and thus represent MMR-dependent mutations.

**Figure 2.**
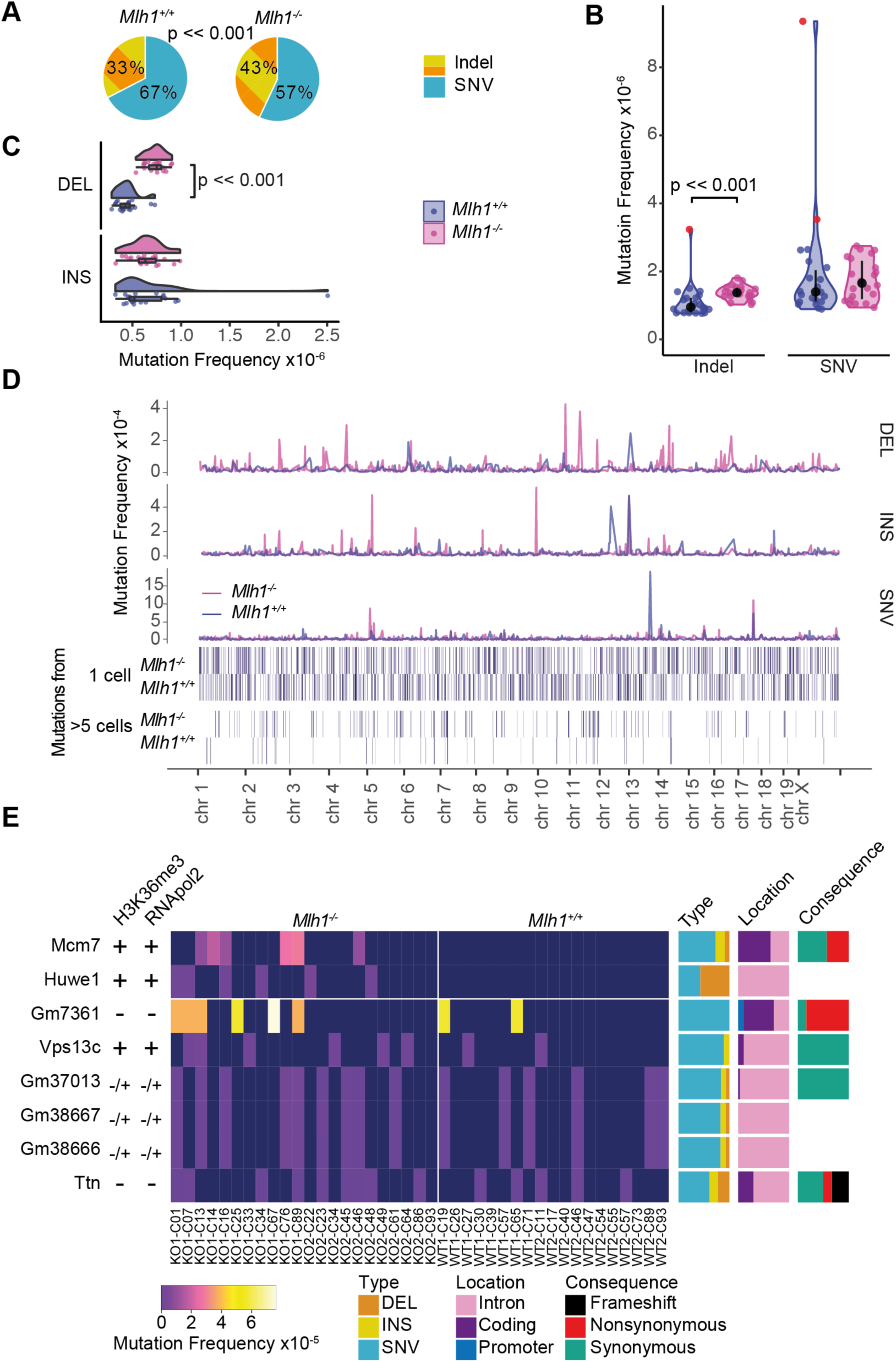
Global and local mutation frequencies in single T cells. (A) *Mlh1*^−/−^ T cells have an increased amount of indels out of total mutations in whole exome compared to *Mlh1*^+/+^ T cells (p << 0.0001, Fisher’s exact test). Global (B) indel and SNV frequencies together with median and interquartile range, and (C) deletion and insertion frequencies in *Mlh1*^+/+^ and *Mlh1*^−/−^ T cells. Mlh1^−/−^ T cells have significantly higher indel, and especially deletion, frequencies than Mlh1^+/+^ T cells (p << 0.001, two-tailed Mann-Whitney U-test). Outlier cells (see methods) are marked with red color in (B). (D) Local mutation frequencies in 1 Mb windows across mouse genome. *Mlh1*^−/−^ T cells have multiple high local mutation peaks originating from only single T cell. (E) Mcm7 and Huwe1 are mutational hotspots in *Mlh1*^−/−^ T cells. Columns are sorted by genotype and cell ID (outliers excluded), rows based on the average mutation frequency. *Mlh1*^+/+^ cells have label WT and *Mlh1*^−/−^ cells have label KO, biological replicates are marked with 1 and 2. Each cell has cell identifier that originates from the Fluidigm C1 plate capture site. Bar plots on the right show proportions of mutation types, locations, and consequences in genes. Left hand side columns show positivity or negativity for RNApol2 and H3K36me3 peaks (**Fig. S3A**).

### Huwe1 and *Mcm7* genes are mutational hotspots in *Mlh1*^−/−^ T cells

*Mlh1*^−/−^ cells provide a unique opportunity to reveal which chromosomal regions represent a particular challenge to the fidelity of the replication machinery, as any errors that are introduced will remain uncorrected by MMR. To identify such regions, we analyzed mutation frequencies in 1 Mb windows across single cell exomes. On a megabase-scale, local mutational frequencies were highly heterogeneous. The majority of the high mutation frequency peaks originated only from single T cells, and mutational hotspot windows shared between individual cells were sparse (Fig. 2D). To establish whether any genes would emerge as MMR-dependent mutational hotspots, we scored all genes for mutations and asked which ones were mutated frequently in *Mlh1*^−/−^ T cells (in more than 5 *Mlh1*^−/−^ cells). Two genes, *Huwe1* and *Mcm7*, stood out with their high mutational frequencies, exclusive to *Mlh1*^−/−^ single cell exomes (Fig. 2E). *Huwe1* encodes an E3 ubiquitin ligase, shown to regulate hematopoietic stem cell self-renewal and proliferation, and commitment to the lymphoid lineage (23). *Mcm7* encodes a component of the MCM2-7 complex that forms the core of the replicative helicase, responsible for unwinding DNA ahead of the replication fork (24). However, only *Mcm7* possessed potentially deleterious mutations in our data set (Fig. 2E). Both genes are positive for RNA polymerase 2 and H3K36me3 in the mouse thymus and expressed from hematopoietic stem cells all the way to thymic T cells (Fig. 2E, Fig. S3A-B).

We then compared the mutational hotspots in *Mlh1*^+/+^ and *Mlh1*^−/−^ normal T cells (this study) and with those in *Mlh1*^−/−^ T cell lymphomas (16). Only one shared mutational hotspot gene was found: *Ttn*, a massive gene with 324 exons, was mutated in both *Mlh1*^−/−^ and *Mlh1*^+/+^ single cell exomes (Fig 2E). We did not identify any mutations in *Ikzf1*, previously reported as a mutational target gene in *Mlh1-*deficient T cell lymphomas (16, 25). Other identified hotspot genes (*Gm7361, Vps13c, Gm37013, Gm38667, Gm38666*) were mutated in both *Mlh1*^−/−^ and *Mlh1*^+/+^ T cells, and thus were not specific for *Mlh1-*deficiency. All except *Vps13c* were negative or inconclusive for the presence of H3K36me3 and RNA polymerase 2, suggesting that these genes are not transcribed in mouse thymus (Fig. 2E, Fig. S3A). *Gm37013, Gm38667* and *Gm38666* are predicted genes and they physically overlap with each other on chromosome 18 (**Fig. S3A**), which explains their identical mutational pattern.

### Insertions and deletions accumulate differently within repeats in *Mlh1*^+/+^ and *Mlh1*^−/−^ T cells

Next, we analyzed the size distribution of detected indels in single cell exomes. *Mlh1*^+/+^ cells had more 1-nucleotide (nt) insertions than deletions, while this difference in *Mlh1*^−/−^ T cells was evened out by increased 1-nt deletions (O.R = 1.794, 95% CI = 1.531-2.101, p = 1.134×10^−13^, Fig. 3A). The same trend for 1-nt insertions as the dominant indel type in *Mlh1*^+/+^ cells was observed in bulk T cell DNA samples from the same mice (**Fig. S4**).

**Figure 3.**
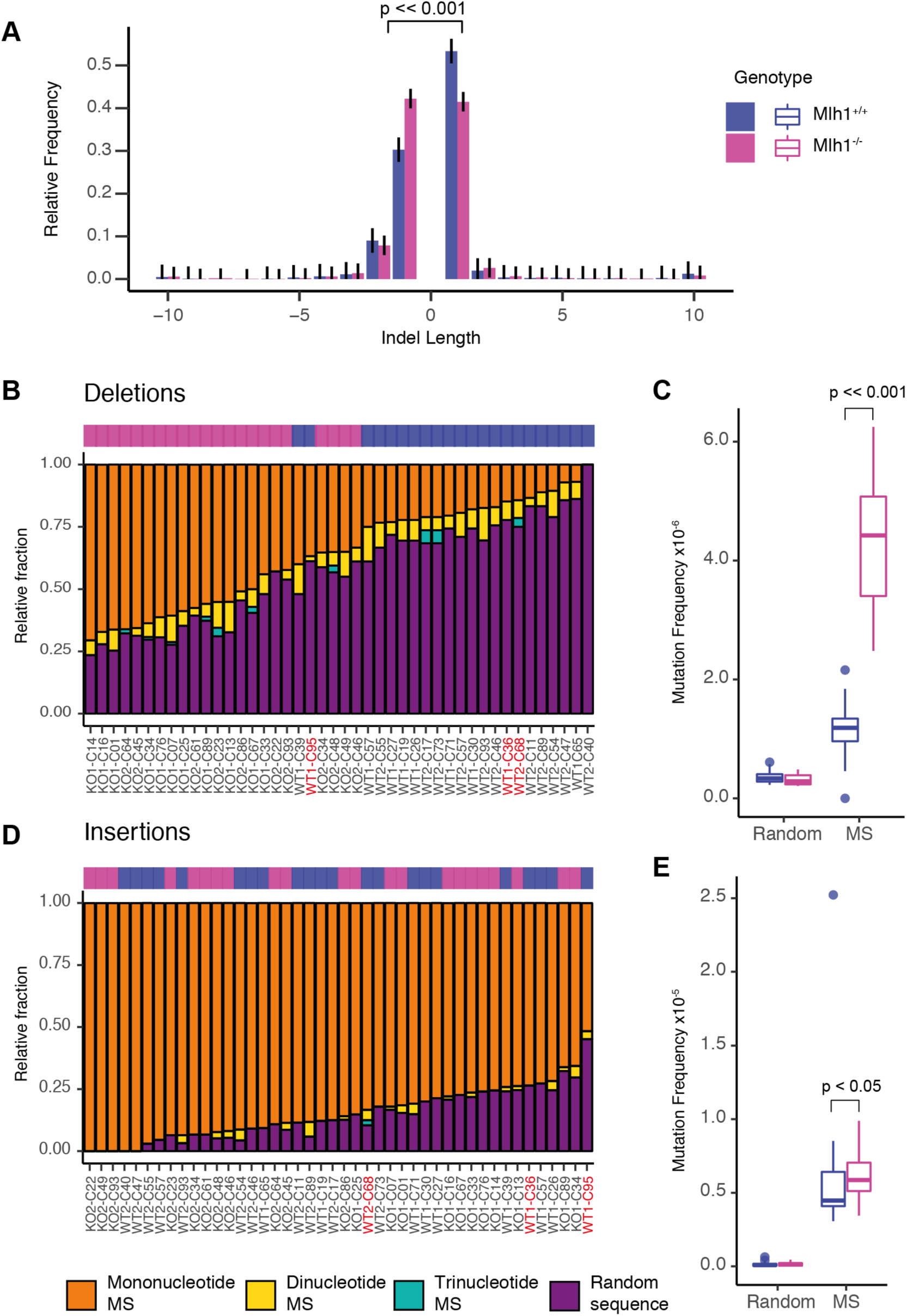
Small deletions report on MMR dependent mutations in mouse T cells. (A) Indel length distribution as relative frequencies with Sison and Glaz 95% multinomial confidence intervals in *Mlh1*^+/+^ and *Mlh1*^−/−^ T cells. *Mlh1*^−/−^ and *Mlh1*^+/+^ cells have different ratios of 1-nt indels (p << 0.001, two-tailed Fisher’s exact test). Indels of length ≥10 bp are binned together. (B) Relative and (C) normalized frequencies of deletions in microsatellites (MS) (mono-, di- and trinucleotide repeats) and in non-microsatellite (random) sequence in single-cell samples. (D) Relative and (E) normalized frequencies of insertions in microsatellites (mono-, di- and trinucleotide repeats) and in non-microsatellite (random) sequence in single-cell samples. Bar plots are ranked by descending mutation fraction within mononucleotide repeats. *Mlh1*^−/−^ cells have a significantly higher deletion frequencies in microsatellites than *Mlh1*^+/+^ (p << 0.001, two-tailed Mann-Whitney U-test). Mutation frequencies are shown as boxplots. Outliers (see methods) are labeled with red in (B) and (D).

We then analyzed the sequence context of the detected indels. As expected, most deletions in *Mlh1*^−/−^ cells occurred at mononucleotide microsatellites, while in *Mlh1*^+/+^ cells, most deletions were found in non-microsatellite sequences (Fig. 3B). When deletion counts were corrected for the number of base pairs of either microsatellite or non-microsatellite sequences, deletion frequencies were higher in microsatellites than in non-microsatellite sequences, regardless of MMR status (Fig. 3C). This underscores the well-documented intrinsic propensity of microsatellites to slippage during replication. As expected, *Mlh1*^−/−^ cells had significantly higher deletion frequencies in microsatellite sequences compared to *Mlh1*^+/+^ cells (p = 9.505×10^−13^, Fig. 3C, Table S1). Insertion frequencies within repeats were more similar between *Mlh1*^−/−^ and *Mlh1*^+/+^ T cells, occurring especially in mononucleotide repeats (Fig. 3D). *Mlh1*^−/−^ cells had slightly higher insertion frequencies in the context of microsatellite sequences (p =0.039, Fig. 3E, Table S1).

### Exons show a decreased burden of MMR-dependent mutations

Exome sequencing, despite its name, not only captures exons, but also exon-adjacent, non-coding regions (Fig. 1) (26). This enabled us to ask whether *de novo* mutations accumulate differently in these two functionally distinct genic regions (exonic versus non-coding) in *Mlh1*^−/−^ and *Mlh1*^+/+^ cells.

No significant difference in SNV frequencies or insertions was observed in either exonic or non-coding regions in *Mlh1*^−/−^ cells compared to *Mlh1*^+/+^ cells (Fig. 4A-B). In contrast, deletions frequencies increased in *Mlh1*^−/−^ cells in non-coding regions compared to *Mlh1*^+/+^ cells (p = 9.94×10^−5^, Fig. 4C, Table S1). Exonic deletion frequencies in *Mlh1*^−/−^ cells did not differ from those observed in *Mlh1*^+/+^ cells (Fig. 4C), indicating that in the absence of functional MMR, the integrity of coding regions is still maintained, likely by purifying selection, as for MMR-deficient tumors by Kim et al., 2013. In conclusion, MMR-dependent mutations increased more in non-coding regions adjacent to exons, as compared to exons themselves.

**Figure 4.**
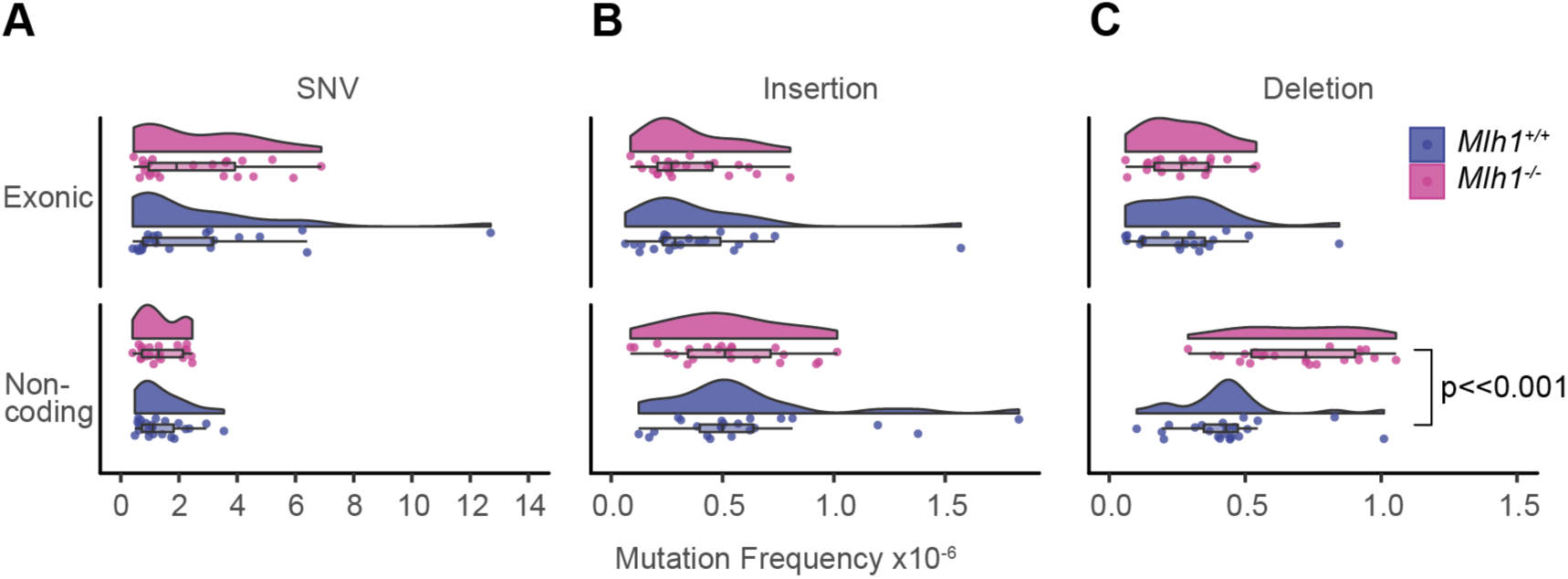
*Mlh1*^−/−^ cells accumulate mutations to non-coding regions of genome. (A) SNV, (B) insertion and (C) deletion frequencies in exonic and non-coding (3’ and 5’ UTRs, promoters, splice sites, introns) regions of the exome in *Mlh1*^+/+^ and *Mlh1*^−/−^ T cells. *Mlh1*^−/−^ T cells have significantly higher frequencies of non-coding deletions (p<<0.001, Two-tailed Mann-Whitney U-test).

### H3K36me3-enriched regions are depleted of MMR-dependent mutations

Results from large tumor data sets strongly indicate that exons have a decreased mutation burden due to H3K36me3-mediated MMR (10), but evidence of this in normal cells and tissues *in vivo* is still lacking. To assess whether replication errors in transcribed genes are buffered by MMR by virtue of their H3K36me3 enrichment, we first analyzed H3K36me3 abundance in RNA polymerase 2 (RNApol2)-positive (RNApol2^+^) and -negative (RNApol2^−^) genes in thymus using publicly available ChIP-seq data (27, 28). Presence of RNA polymerase 2 in the promoter region is a strong indicator of transcriptional activity (29). H3K36me3 levels in RNApol2^+^ regions were higher than in RNApol2^−^ regions and peaked at the centers of these regions (Fig. 5A), confirming that H3K36me3 is associated with transcriptional activity also in mouse thymus. However, not all RNApol2^+^ genes were positive for H3K36me3. Approximately 65% of RNApol2^+^ genes were also positive for H3K36me3, whereas 80% of H3K36me3 positive (H3K36me3^+^) genes were positive for RNApol2 (Fig. 5B).

**Fig 5.**
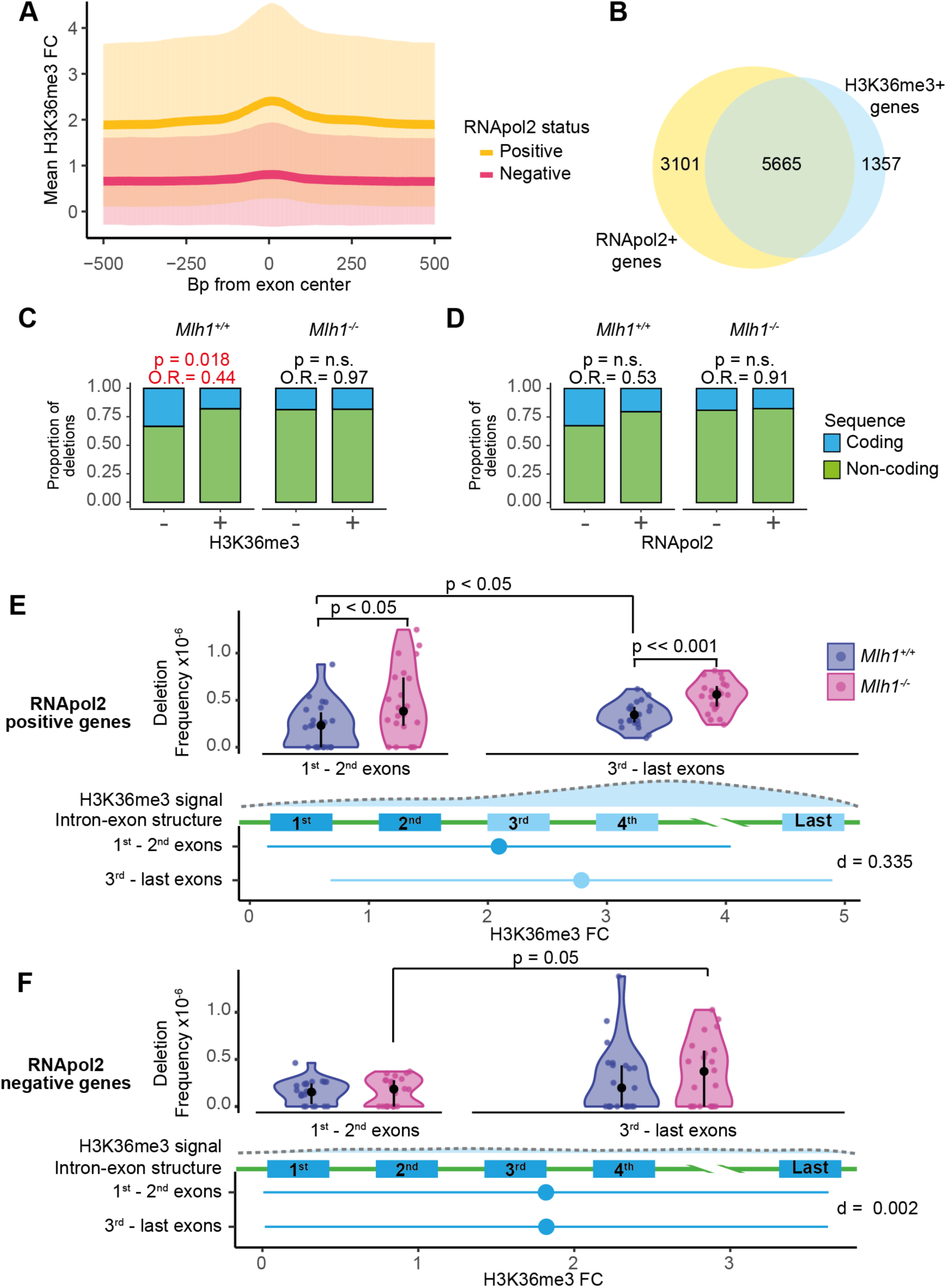
H3K36me3 reduces the amount of MMR-dependent mutations in exons. (A) H3K36me3 fold change (FC) (mean ± s.d.) in 1000 bp window around of exon centers in RNApol2 positive and negative genes. (B) Venn diagram of RNApol2 positive (+) and H3K36me3 positive (+) gene counts. Proportions of small deletions and insertions in genes positive or negative for (C) H3K36me3 and (D) RNApol2. Coding regions in genes positive for H3K36me3 have less deletions relative to silent genes in *Mlh1*^+/+^ cells (p = 0.018, O.R. = 0.44, two-tailed Fisher’s exact test), but not in *Mlh1*^−/−^ cells. Deletion frequencies in 1^st^ to 2^nd^ exons (5’ exons) and 3^rd^ to last exons (3’ exons) in RNApol2 (E) positive and (F) negative genes. In RNApol2-positive genes, *Mlh1*^−/−^ cells have higher deletion frequency especially in 3^rd^ to last exons (high H3K36me3) than *Mlh1*^+/+^ cells, and to lesser degree, in the 1^st^ to 2^nd^ exons (low H3K36me3). First panel shows the deletion frequencies together with median and interquartile range in *Mlh1*^+/+^ and *Mlh1*^−/−^ cells. Second panel shows a schematic of H3K36me3 enrichment along a gene. Third panel shows a schematic of a gene structure. Fourth panel shows H3K36me3 signal as mean ± s.d. of FC in 1^st^ to 2^nd^ exons and 3^rd^ to last exons together with effect size as Cohen’s d with Bessel’s correction. Deletion frequencies were tested using two-tailed Mann-Whitney U-test.

We analyzed how small deletions (that is, MMR-dependent mutations) were distributed to exons and non-coding regions based on either RNApol2 or H3K36me3 status of genes. The proportion of exonic deletions over non-coding deletions was decreased in H3K36me3^+^ genes compared to H3K36me3-negative (H3K36me3^−^) genes in *Mlh1*^+/+^ (p = 0.018, OR = 0.44, 95% CI = 0.198-0.906), but not in *Mlh1*^−/−^ T cells (p = 1, OR = 0.972, 95% CI = 0.542-1.694, Fig. 5C). Lower exonic deletion burden in RNApol2^+^ genes was also observed in *Mlh1*^+/+^ cells (Fig. 5D), similar to H3K36me3^+^ genes (p = 0.062, OR = 0.528, 95% C1 = 0.250-1.060, Fig. 5C). The similar trends are not surprising, given the overlap between RNApol2^+^ and H3K36me3^+^ genes (Fig. 5B). These results strongly support H3K36me3-guided, MMR-dependent protection of exons against genetic alterations.

The H3K36me3 mark is less abundant in 5’ exons, compared to 3’ exons of genes (6, 10). To test whether local H3K36me3 levels affect the intra-genic distribution of mutations within genes, we compared deletion frequencies in 1^st^ and 2^nd^ exons (from here on referred to as 5’ exons) with those in 3^rd^ to last exons (from here on referred to as 3’ exons), both in RNApol2^+^ and RNApol2^−^ genes. In RNApol2^+^ genes, H3K36me3 signal increased in 3’ exons compared to 5’ exons (d = 0.335, Fig. 5E), whereas in RNApol2^−^ genes, there was no difference in H3K36me3 levels between 3’ and 5’ exons (d = 0.002, Fig. 5F, Table S1). In RNApol2^+^ genes, *Mlh1*^−/−^ cells had higher deletion frequencies in 3’ exons (high in H3K36me3) compared to *Mlh1*^+/+^ cells (p = 4.57×10^−5^, Fig. 5E, Table S1). In 5’ exons (low in H3K36me3), the difference in deletion frequencies between *Mlh1*^−/−^ and *Mlh1*^+/+^ was smaller, yet significant (p = 0.016, Fig. 5E, Table S1). *Mlh1*^+/+^ cells also had somewhat increased deletion frequencies in the 3’ exons compared to 5’ exons (p = 0.020, Fig. 5E, Table S1). Sequencing coverage was similar between samples with or without mutations in the analyzed exons, except in the 5’ exons in RNApol2^+^ regions in *Mlh1*^+/+^ cells (p = 0.04, **Fig. S5**). Taken together, these results suggest that 3’ exons in transcriptionally active genes are more prone to acquiring mutations compared to 5’ exons, and that this effect is tempered by H3K36me3-guided MMR. No difference was observed in the deletion frequencies between *Mlh1*^+/+^ and *Mlh1*^−/−^ cells in the RNApol2^−^ genes in 5’ exons (p = 0.539) or 3’ exons (p = 0.296, Fig. 5F, Table S1). *Mlh1*^−/−^ cells, however, showed a small difference between deletion frequencies in 5’ exons and 3’ exons (p = 0.049, Fig. 5F, Table S1). H3K36me3^−^ exons in RNApol2^−^ genes accumulated mutations in similar frequencies in both *Mlh1*^+/+^ and *Mlh1*^−/−^ cells. We interpret this to mean that the MMR machinery does not operate efficiently in these regions even in wildtype cells. RNApol2^+^, but not RNApol2^−^ genes showed genotype-dependent spatial variability in deletion frequencies, thus transcriptional activity appears to affect accumulation and/or repair of replication errors.

## DISCUSSION

Using single-cell exome sequencing of mouse thymic T cells, we uncovered how the exome-wide mutational landscape is shaped *in vivo* by replication errors and by MMR-mediated error correction. We further provide evidence for transcription-associated replication errors and H3K36me3-guided MMR at 3’ exons of genes.

We show that scWES is a sensitive approach for unraveling signatures of replication errors and MMR activity. This is highlighted by the fact that we detected a substantial increase of deletions in *Mlh1*^−/−^ T cells, and found evidence of insertional bias in *Mlh1*^+/+^ T cells.

DNA polymerases tend to create more deletions than insertions, especially in repeat sequences (30–35). In the absence of MMR (which is the situation in *Mlh1*^−/−^ cells), one would expect to directly detect replication errors. Indeed, we observed a significant increase of small deletions in *Mlh1*^−/−^ cells compared to *Mlh1*^+/+^ cells, as expected given the deletion bias of DNA polymerases. Taken together, we conclude that deletions reliably report of the replication errors that would otherwise be repaired by MMR.

In addition, we found that *Mlh1*^+/+^ cells had more insertions than deletions. Increase in 1-nt insertions rather than deletions in *Mlh1*^+/+^ cells has also been observed at unstable microsatellite loci in other MMR-proficient normal mouse tissues (36). Our findings are in line with the previously reported bias for MMR to correct deletions more efficiently than insertions, thereby creating an insertional bias at microsatellites (37).

*Mlh1-*deficient cells lack MMR activity and accumulate replication-induced errors with every cell division. Developing lymphocytes are particularly susceptible to replication errors because they undergo multiple rounds of proliferative expansions during development and maturation. Comparison of mutational frequencies in *Mlh1*^−/−^ versus *Mlh1*^+/+^ T cell exomes revealed two hotspots for replication errors, *Huwe1* and *Mcm7*genes. Because these genes appear vulnerable for replication errors, we propose, that over time, in *Mlh1*-deficient cells damaging mutations will likely emerge. Indeed, mutations in *Huwe1* and *Mcm7* have been reported in a subset of *Mlh1*-deficient murine T cell lymphomas (16). The propensity of *Mcm7*, coding for an integral component of the replication machinery, to acquire deleterious mutations in MMR-deficient cells (Figure 2E) conceivably can accelerate the accumulation of replication-associated errors, thereby adding insult to injury.

Both *Huwe1* and *Mcm7* are expressed in the T lymphocyte lineage and required for lymphocyte development. Shielding them from permanent mutations is likely important for cellular homeostasis and normal development, and *Huwe1* and *Mcm7* were in fact devoid of mutations in *Mlh1*^+/+^ T cells. In the face of frequent replication errors, how is efficient targeting of MMR to these regions ensured in wildtype cells? Both *Huwe1* and *Mcm7* were enriched for H3K36me3 in the mouse thymus, and H3K36me3-mediated MMR has been shown to protect actively transcribed genes (11). Thus, H3K36me3-mediated recruitment of MMR machinery to these genes provides an explanation for efficient error correction in wildtype cells; in the absence of MMR, H3K36me3 no longer has a protective effect.

Also, on a single cell resolution, the protective effect of H3K36me3-mediated MMR on active genes appears to hold true more globally. On the whole-exome level, MMR-dependent mutation frequencies in wildtype cells were lower especially in H3K36me3-enriched exons when compared to *Mlh1*^−/−^ cells. Our results indicate that H3K36me3-mediated MMR conserves the integrity of active genes in normal tissues *in vivo*, similarly as shown previously for tumors and cell lines (8, 10, 11).

Moreover, we show evidence that 3’ ends of actively transcribed genes are more prone to replication-associated errors and that more efficient recruitment of MMR via H3K36me3 protects these regions and ensures that most of these errors do not become permanent mutations. Head-on collisions of the replication and transcription machineries can cause indels and base-substitutions, and especially increase the deletion burden within 3’ ends (and to a lesser degree 5’ ends) of genes under active transcription (38). Moreover, SNVs accumulate more to 3’ UTRs than to 5’ UTRs in aging B lymphocytes (19), supporting the observation that 3’ regions are in fact more prone to mutations. Efficient recruitment of the MMR machinery via H3K36me3 can shield against replication-induced errors specifically in transcribed genes, whose integrity is particularly important.

## CONCLUSIONS

Here, we delineate the mutational landscape of T cells shaped by the status of DNA repair (functional vs impaired), dissected at the single-cell level in the context of H3K36me3. We provide evidence that in normal T cells, MMR preferentially protects genes, and in particular H3K36me3-positive 3’ exons transcribed in T cell lineage, against accumulation of *de novo* mutations. Taken together, our results suggest an attractive concept of thrifty MMR targeting, where genes critical for the development of a given cell type and under mutational stress due to active transcription are preferentially shielded from deleterious mutations.

## MATERIALS AND METHODS

### Mice

Two female *Mlh1*^−/−^ (13) and two of their *Mlh1*^+/+^ female littermates, age 12 weeks, were used for the single-cell whole exome sequencing study.

### Enrichment of thymic T cells

Mice were euthanized by carbon dioxide inhalation, followed by cervical dislocation. Thymi were collected in ice-cold DMEM (Gibco cat: 11960-044) and visually inspected for any macroscopic anomalies. Whole thymi were homogenized for an enrichment of naïve T cells using a commercially available kit according to manufacturer’s instructions (Invitrogen, cat:11413D).

### Single-cell capture and whole genome amplification

Enriched T cells were prepared for single-cell capture and whole-genome amplification in Fluidigm C1 system according to manufacturer’s protocol (Fluidigm cat: 100-7357). Single T cells were captured using an IFC 5-10 μm capture plate (Fluidigm cat: 100-5762) and imaged using Nikon Eclipse Ti-E microscope with Hamamatsu Flash 4.0 V2 scientific CMOS detector. After confirming the capture by microscopy, cell lysis and whole-genome amplification steps were carried out in Fluidigm C1 system using illustra GenomiPhi V2 DNA Amplification Kit (GE Healthcare Life Sciences cat: 25-6600-30). DNA concentrations of amplified single-cell genomes were determined using either a Qubit dsDNA HS Assay kit (Invitrogen cat:Q32854) with Qubit Fluorometer (1.27) or QuantiFluor dsDNA System (Promega cat:E2670) with Quantus Fluorometer (2.24). Fragment size and integrity of amplified single-cell genomes were analyzed using Bioanalyzer High Sensitivity DNA Assay (Agilent) with Agilent Bioanalyzer 2100 (2100 Expert B.02.08.S648 SR3) or TapeStation Genomic DNA ScreenTape (Agilent) with TapeStation 4200 (TapeStation Analysis Software A.02.021 SR1) at the Biomedicum Functional Genomics Unit, Helsinki. Samples with the highest density of fragments around ~10 kb were chosen for sequencing based on visual inspection of the fragment size distributions.

### Library preparation and sequencing

Agilent SureSelectXT Mouse All Exon 49.6Mb capture was used for exome enrichment and to prepare multiplexed libraries for Illumina. Samples were sequenced using Illumina NextSeq 500 with mid output reagents as paired-end 150 bp reads. In total, we sequenced 56 single T cell exomes in three batches, each batch consisting of single-cell samples with a genotype-matched bulk DNA sample (= whole genome amplified cell suspension, n=3/genotype, biological replicates 1 and 2, and technical replicate for biological replicate 1). Sequencing was performed by the Biomedicum Functional Genomics Unit, Helsinki.

### Sequence alignment

Sequence alignment and variant calling workflow was adapted from Leung et al. (39). Paired-end reads were aligned to the Dec. 2011 (GRCm38/mm10) assembly of the mouse genome using bowtie2 (2.3.4) (40) with --local mode. Aligned reads were then sorted, merged, and marked for duplicates using SAMtools (1.4) (41) and Picard (2.13.2) (42). Reads were re-aligned around indels using GATK (3.8-0-ge9d806836) (43), followed by removal of reads with low mapping quality (MQ < 40) using SAMtools. Sequencing metrics (average depth and coverage) were calculated using SAMtools, BEDtools (2.26.0) (44) and R (3.5.0). Samples that had coverage less than 50% at depth ≥1X were excluded from subsequent analyses (**Fig. S1B**).

### Variant calling and filtering

Variants within the exome capture region + 100 bp interval padding were called using GATK HaplotypeCaller in -ERC GVCF mode, followed by joint calling with GenotypeGVCFs. Samples (single-cell and bulk DNA) from the same genotype (*Mlh1*^+/+^ or *Mlh1*^−/−^) were analyzed together. Variant score recalibration was done separately to indels and SNVs using GATK SelectVariants and VariantRecalibration and applied at 99.0 sensitivity level using ApplyRecalibration. Variant sets used to build the recalibration model for SNVs were dbSNP (build 150) (45), Mouse Genomes Project SNP Release Version 5 (46), and bulk SNV set (see below), and for indels, dbSNP (build 150), Mouse Genomes Project indel Release Version 5, and bulk indel set (see below). After variant score recalibration, all variants that had genotype quality <20, depth <6 and heterozygous genotypes allelic depth <0.333 were filtered out. Clustered SNVs (>3 SNVs / 10 bp) were filtered out to eliminate false positive SNVs caused by poor alignment around indels. Variants found in both *Mlh1*^+/+^ and *Mlh1*^−/−^ samples (germline mutations), homozygous mutations (insufficient whole-genome amplification) and variants found in the 129P2 OlaHsd strain were excluded from all subsequent analyses (mice with disrupted *Mlh1* were originally created using 129/Ola derived embryonic stem cells that were injected to C57BL/6 mice (13)). Filtering was done using GATK VariantFiltration, Picard FilterVcf, and R package *VariantAnnotation* (1.26.1) (47).

### High confidence bulk indel and SNV training set construction

High confidence bulk DNA SNV and indel training sets for variant score recalibration were constructed from the raw variants discovered in bulk DNA samples (both *Mlh1*^+/+^ and *Mlh1*^−/−^) by including the variants that passed the following filters: ReadPosRankSum > −1.9, QD > 5.0, SOR > 1.5 for indels and SNVs, and for SNVs only: MQRankSum > −1.9. Variants that did not have a genotype (= insufficient sequencing coverage) across all bulk samples (n=3/genotype) were removed from the reference bulk set.

### Mutation annotation

Mutations were annotated (gene, genic location, mutation consequence) using R package *VariantAnnotation* function *locateVariants* with *AllVariants* option and *predictCoding*. The UCSC KnownGene track from *TxDb.Mmusculus.UCSC.mm10.knownGene* (3.4.0) was used as the gene model. We considered mutations that fall within CDS regions to be exonic, and those that fall within 5’ untranslated region (UTR), 3’ UTR, splice site, intron or promoter to be non-coding. For analysis of exonic and non-coding indels (Fig. 4A-C and Fig. 5C-D), we included mutations in genes with only one transcript to avoid having multiple locations within one gene for one mutation. In the mutation hotspot analysis (Fig. 2E), all possible transcript variants were analyzed.

### Regions with transcriptional activity and enriched with H3K36me3 in mouse exome

RNApol2 (ENCFF119XEH) and H3K36me3 (ENCFF853BYO) ChIP-seq peak coordinates for mouse thymus were downloaded as BED files from ENCODE (27, 28). We used UCSC knownGene track to define the genomic coordinates of genes. Genes that overlapped or were within 100 bp of the ChIP-seq peak coordinates were defined positive for that feature. Genes positive for H3K36me3 or RNApol2 peaks were defined separately.

### H3K36me3 signal in genes

H3K36me3 data (ENCFF287DIJ) for mouse thymus was downloaded as a BigWig file containing fold-change (FC) of ChIP reads over background reads from ENCODE (27, 28). Mean H3K36me3 FC ± standard deviation (s.d.) in each position (meaning, each *base* gets a mean H3K36me3 FC value) 500 bases up- and downstream from the exome capture centers was calculated for RNApol2-positive and -negative genes. Mean H3K36me3 FC ± s.d. in 5’ and 3’ exons (meaning, each *region* gets a mean H3K36me3 FC value) were calculated for RNApol2-positive and -negative genes.

### Microsatellites in mouse exome

Mono-, di-, and trinucleotide repeats in mouse exome were detected using STR-FM (Galaxy version 1.0.0) (48) in Galaxy at usegalaxy.org (49). R package *BSgenome.Mmusculus.UCSC.mm10* (1.4.0) was used to convert BED file containing genomic coordinates of variant call regions into FASTA format. Mono-, di-, and trinucleotide repeats were detected from the FASTA file in separate runs using motif sizes 1, 2, and 3, no partial motifs allowed, and minimum repeat unit counts were 4 (minimum length 4 bp) in mononucleotide repeat detection and 3 in dinucleotide (minimum length 6 bp) and trinucleotide (minimum length 9 bp) repeat detections. Non-microsatellite associated regions were defined as those that were not defined as mono-, di-nor trinucleotide repeats.

### Microsatellite associated indels in single-cells

Sequence 100 bp up- and downstream of detected indel start coordinates were extracted from the mouse reference genome mm10 (*BSgenome.Mmusculus.UCSC.mm10*) in FASTA format and analyzed for mono-, di- and trinucleotide repeats as described above. Indels were marked microsatellite-associated if the indel start coordinate and microsatellite start coordinate were the same. Indels found not to be within mono-, di- or trinucleotide repeat were labelled as non-microsatellite associated (random) indels.

### Mutation frequencies in single T cells

Global indel and SNV frequencies in the variant call region were calculated for each single-cell and reported as mutations/base. Mutation frequency was calculated as: *frq =n/(cov*2)*, where *n* is the number of mutations, *cov* is the number of high-quality base pairs (MQ > 40, DP > 6). Similarly, frequencies in different genomic regions (exonic, non-coding, microsatellites, 3’ exons, 5’ exons) were calculated by first counting the number of mutations in each region and dividing it by the coverage of that particular region.

### Mutation frequencies in 1 Mb windows

Local mutation frequencies in 1 Mb windows were calculated by first dividing the genome into 1 Mb windows, then calculating the coverage of variant call region (exome capture + 100 bp padding) in each window. Next, the number of SNVs, deletions, and insertions per genotype (*Mlh1*^+/+^ or *Mlh1*^−/−^) was counted in each window. Mutation frequency for *Mlh1*^+/+^ and *Mlh1*^−/−^ groups was then calculated by dividing the number of observed mutations in each window by the coverage (*cov*2*) of variant call region in that window.

### Mutation hotspot analysis

We analyzed all genes for mutations in *Mlh1*^+/+^ and *Mlh1*^−/−^ T cells. For each sample, we counted the number of mutations per gene. These numbers were then normalized by the coverage (*cov*2*) of the gene in each sample. A gene was considered to be a hotspot if it was mutated in more than 5 *Mlh1*^−/−^ T cells.

### Outlier cells in single-cell samples

Cells that had indel or SNV frequency higher or lower than 1.5 * interquartile range in matching genotype were labelled as outliers and removed from all the subsequent statistical test. Outliers are shown in the plots, unless mentioned otherwise, and indicated in Figs. 2B, 3B and 3D.

### MMR dependent mutation frequencies in 5’ and 3’ exons

To analyze mutation frequencies and H3K36me3 signal in 5’ exons (1^st^ to 2^nd^ exons) and 3’ exons (3^rd^ to last exons), we took UCSC knownGene transcripts, excluded genes that overlap each other, and collapsed transcripts gene-wise to create one exon-intron-structure for each gene. 100 bp padding was added to each exon. Only genes with 4 or more exons were considered and exons 1-2 were marked as 5’ exons ad exons 3-last were marked as 3’ exons. Genes that were in or within 100 bp of RNApol2 peak coordinates were marked as RNApol2 positive. Number of deletions in 5’ and 3’ exons in each single-cell were counted and then divided by the coverage (*cov*2*) of either 3’ or 5’ exons in that single-cell sample.

### General R packages

R version 3.5.0 was used to analyze the data. *VariantAnnotation* package was used for VCF file manipulation, *rtracklayer* (1.40.3) (50) package for reading BED and BigWig files, and *GenomicRanges* (1.32.6) (51) package for handling genomic coordinates in R environment. Figures and general data manipulation were done using *ggplot2* (3.00.0), *gplots* (3.00.1), *Gviz* (1.24.0), *grid* (3.5.0), *viridis* (0.5.1), *dplyr* (0.7.6), *plyr* (1.8.4), *reshape2* (1.4.3), *tidyr* (0.8.2), *VennDiagram* (1.6.20), and *Hmisc* (4.1-1).

### Statistical analysis

All tests were calculated using 22 *Mlh1*^−/−^ T cells and 19 *Mlh1*^+/+^ T cells, except in the Fig. 2A, where all single cell samples were included (22 *Mlh1*^−/−^ T cells and 22 *Mlh1*^+/+^ T cells). All mutation frequencies are reported as median (mdn) and interquartile range (iqr) (**Table S1**) and tested using two-tailed Mann-Whitney U test (*wilcox.test*). P-values for mutation counts (indels and SNVs (Fig. 2A), 1-nt indels in *Mlh1*^+/+^ and *Mlh1*^−/−^ cells (Fig. 3A), mutations in exonic vs non-coding regions in active and silent genes (Fig. 5C-D)) were calculated using two-tailed Fisher’s exact test (*fisher.test*) and reported with odds ratio (O.R., ratio of ratios) and 95% condifence intervals (CI). O.R. values close to 1 indicate no difference in the ratios. Differences were determined statistically significant at a confidence level of 95%. Errors bars shown in Fig. 3A are Sison and Glaz 95% multinomial confidence intervals from R package *DescTools* (0.99.25). Effect size reported for H3K36me3 signal in Fig. 5E-F was calculated using Cohen’s d with Bessel’s correction, implemented in R. Cohen’s d values closer to 0 indicate smaller difference between two group means

## DECLARATIONS

### Ethics approval and consent to participate

All animal experiments were performed following national and institutional guidelines (the National Animal Experiment Board in Finland and the Laboratory Animal Centre of the University of Helsinki) under animal license number ESAVI/1253/04.10.07/2016.

### Consent for publication

Not applicable.

### Availability of data and materials

Single-cell exome sequencing data generated and analyzed during the current study are available as raw reads in FASTQ format in the SRA repository, under accession number <u>PRJNA575619</u>. Publicly available H3K36me3 (<u>ENCFF853BYO</u> and <u>ENCFF287DIJ</u>) and RNApol2 (<u>ENCFF119XEH</u>) ChIPSeq data can be found from ENCODE (https://www.encodeproject.org) database.

### Authors’ contributions

E.A. performed and designed the experiments, performed data analysis, interpreted the results and wrote the manuscript. D.D. designed and performed initial experiments, supervised data analysis and interpretation, and wrote the manuscript. L.K. conceived and designed the study, supervised the experiments, data analysis and interpretation, acquired funding, coordinated the project and wrote the manuscript. All authors read and approved the manuscript.

## Acknowledgements

We are grateful to Fran Supek, Esa Pitkänen, Niko Välimäki and Julia Casado for discussions and advice. We wish to acknowledge CSC – IT Center for Science, Finland for computing resources, the Functional Genomics Unit (University of Helsinki) for sequencing services, Minna Nyström (University of Helsinki) for providing mice, Jussi Taipale and Anna Vähärautio for access to Fluidigm C1 system, and Kul Shanker Shrestha and Minna Tuominen for technical assistance. Assistance was also provided by Laboratory Animal Center, and Biomedicum Imaging Unit at University of Helsinki and Palo Alto Veterans Institute for Research (PAVIR) FACS Core.

