## Supplementary Data for "Single-cell sequencing of mouse thymocytes reveals mutational landscape shaped by replication errors, mismatch repair and H3K36me3"

### SUPPLEMENTARY MATERIALS AND METHODS

#### Analysis of thymic cell populations

Whole thymi of 4 *Mlh1*$^{-/-}$ and 4 *Mlh1*$^{+/+}$ mice were dissected and homogenized in HBSS + 2% FBS + 2 mM EDTA. Cells were frozen in HBSS + 10% DMSO + 40% FBS and stored at -80°C before analysis. Cells were thawed and transferred to warm HBSS + 2% FBS + 2 mM EDTA and washed twice. Live/dead staining was done for 15 minutes in +4°C with AmCyan (Invitrogen) 1:1000, followed by washing with HBSS + 2% FBS + 2 mM EDTA. Cells were stained for 45 minutes at +4°C for APC-TCRγ/δ (clone GL3, BioLegend) (1:400 dilution), AF780-TCRβ (clone H57-597, BD Bioscience) (1:200 dilution), BV711-CD4 (clone RM4-5, BD Biosciences) (1:200 dilution), eVolve655-CD8α (clone 53-6.7, eBioscience) (1:200 dilution), eFluor450-CD3 (clone 17A2, eBioscience) (1:150 dilution), PE-Cy7-NK1.1 (clone PK136, BD Bioscience) (1:200 dilution), BUV737-CD45.2 (clone 104, BD Horizon) (1:100 dilution) with Fc block CD16/32 (Invitrogen) (1:100 dilution) and rat serum (1:10 dilution). Cells were then washed with HBSS + 2% FBS + 2 mM EDTA, and resuspended to PBS. Cells were analyzed by flow cytometry with BD LSRFortessa and results were further analyzed using FlowJo (10.5.3).

#### *Huwe1* and *Mcm7* gene expression in T cell developmental sequence

Single-cell RNAseq gene expression data for hematopoietic stem cells (HSC), Slamf1 negative (-) and Slamf1 positive (+) multipotent progenitors (MPP), common lymphoid progenitors (CLP), pro T cells, immature T cells, and T cells for mm10 were downloaded from

UCSC (http://genome.ucsc.edu/) as txt file tables containing sample IDs, category annotations and TMP values (1).

#### Statistical testing

Percentages of different thymic cell types were tested using an implementation of two-sided permutation test that compares the difference of means to simulated distribution (2). Number of permutations used was 1000.


### SUPPLEMENTARY ACKNOWLEDGEMENTS

We would like to thank Palo Alto Veterans Institute for Research (PAVIR) FACS Core for access to LSRFortessa.


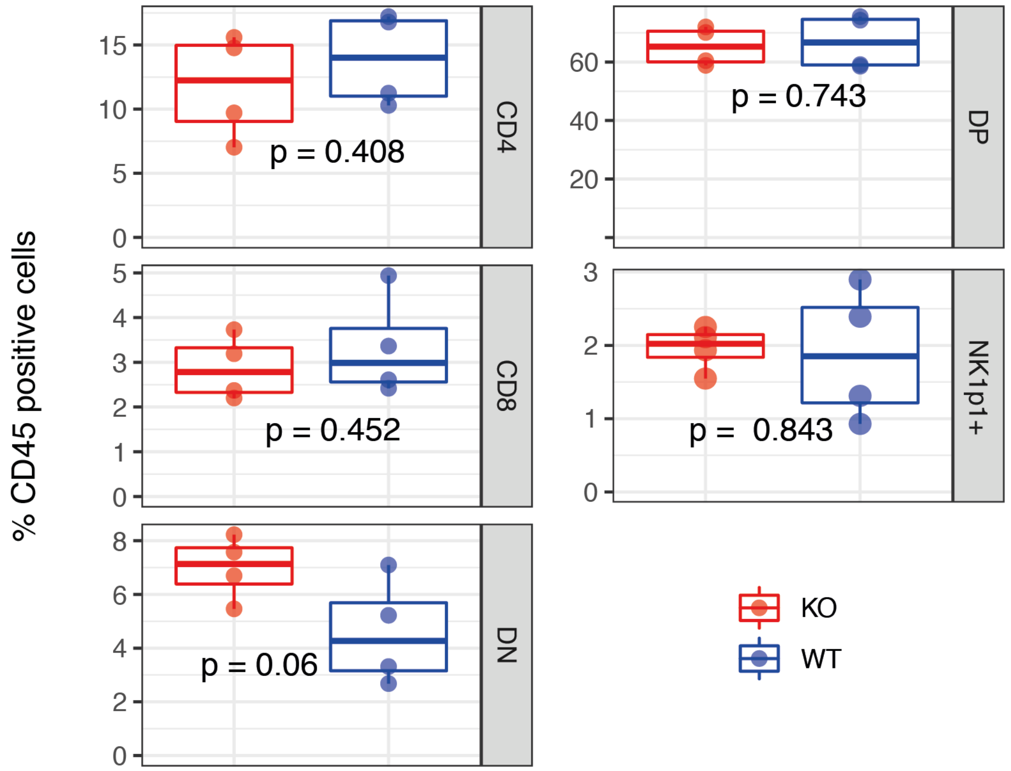


**Figure S1** ***Mlh1***$^{+/+}$ **and** ***Mlh1***$^{-/-}$ **thymic lymphocytes**

CD4-CD8- double negative (DN), CD4+CD8+ double positive (DP), CD8+ single positive (CD8), CD4+ single positive (CD4), and TCRγδ T cells from *Mlh1*$^{+/+}$ (n=4) and *Mlh1*$^{-/-}$ (n=4) whole thymus. No significant difference was observed in different cell populations when comparing *Mlh1*$^{+/+}$ and *Mlh1*$^{-/-}$ mice. P values are two-sided values from permutation test (see Supplementary Methods).

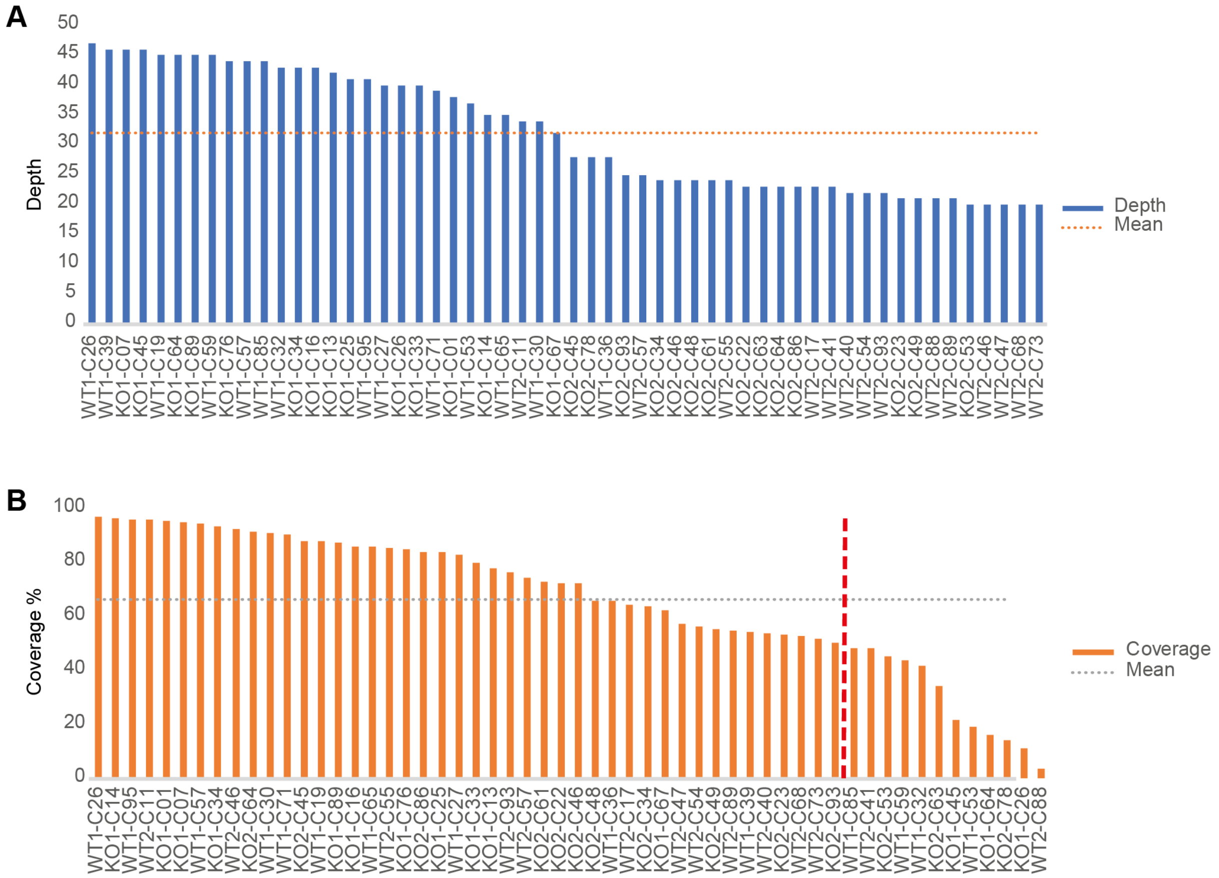


**Figure S2 Average depth and coverage of sequenced single T cell exomes**

(A) Average depth and (B) average coverage at depth ≥ 1 X in single T cells (n = 56). Dashed horizontal line marks mean depth or coverage across all sequenced T cells. Samples to the right of the vertical red dashed line in (B) were excluded from the mutational analysis.

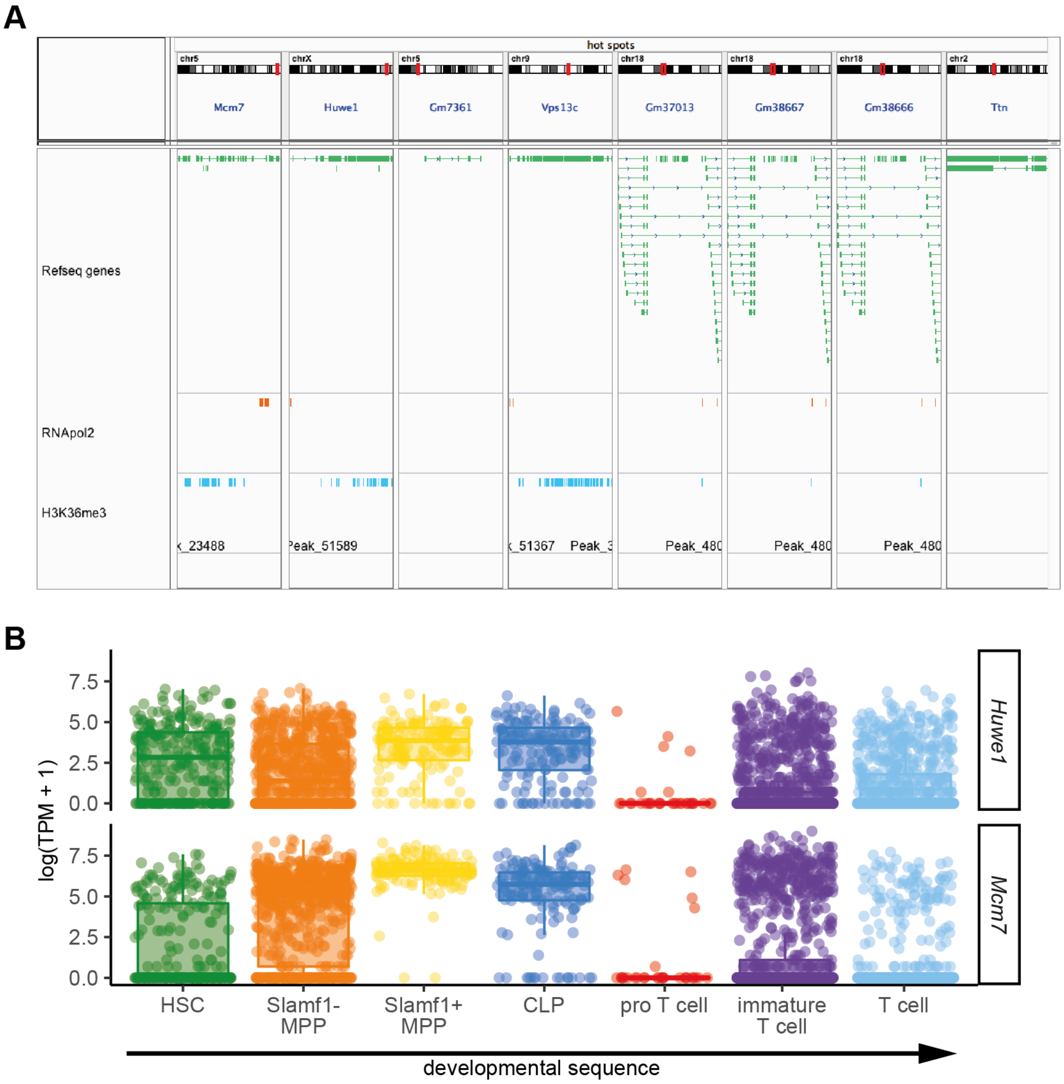


**Figure S3 H3K36me3 enrichment and transcriptional activity in mutational hotspot genes**

(A) RNA polymerase 2 and H3K36me3 ChIP-seq peak locations in mutational hotspot genes. *Mcm7*, *Huwe1* and *Vps13c* are positive (+) for, and *Gm7361* and *Ttn* are negative (-) for both RNA polymerase 2 and H3K36me3. *Gm37013*, *Gm68667* and *Gm38666* are inconclusive (-/+), because of the weak ChIP-seq signal. (B) Single-cell gene expression of *Mcm7* and *Huwe1* in developing T-lymphocytes. *Mcm7* and *Huwe1* are expressed from hematopoietic stem cells to naïve thymic T cells.

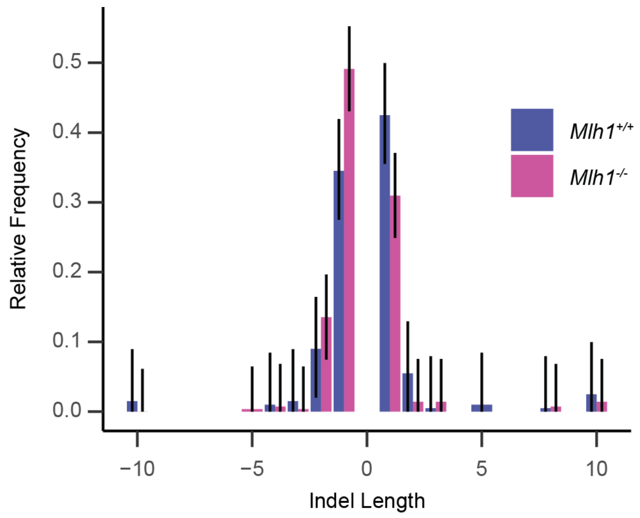


**Figure S4 Indel length distribution in bulk DNA samples**

Indel length distribution as relative frequencies with Sison and Glaz 95% multinomial confidence intervals of *Mlh1*$^{+/+}$ (n = 3) and *Mlh1*$^{-/-}$ (n = 3) bulk DNA samples, shown as relative frequencies. Bulk samples show the same trend in 1-nt indel bias as seen in single-cell samples. Technical replicates: 3, biological replicates: 2.

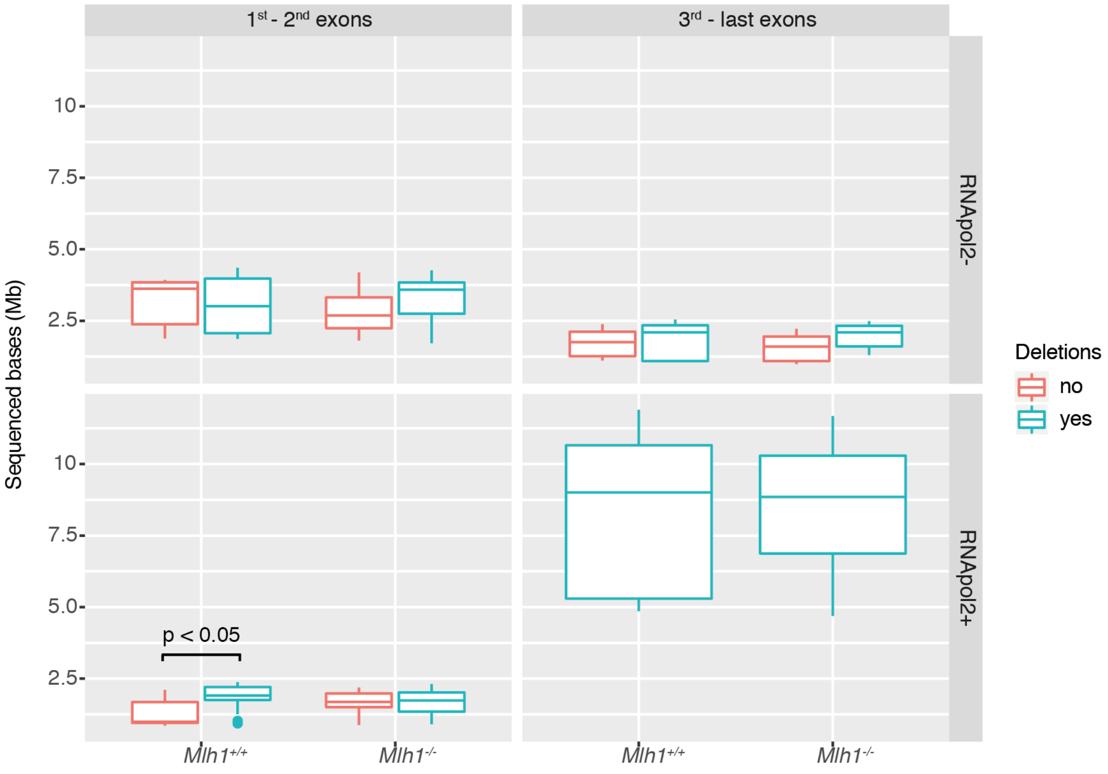


**Figure S5. Sequenced bases in 1st – 2nd exons and 3rd – last exons in RNApol2-negative (top panel) and -positive (bottom panel) genes.** Number of sequenced bases shown as boxplots in 22 *Mlh1*$^{+/+}$ (three outlier cells included, see Methods) and 22 *Mlh1*$^{-/-}$ cells. A significant difference in sequenced bases within different exonic regions between samples with and without deletions was seen only in RNApol2-positive 1st to 2nd exons in *Mlh1*$^{+/+}$ cells ($p = 0.041$, two-tailed Mann-Whitney U-test, n= 19 for *Mlh1*$^{+/+}$, n= 22 for *Mlh1*$^{-/-}$, outlier cells excluded – see Methods).

**Table S1. Mutation frequencies in *Mlh1*$^{+/+}$ and *Mlh1*$^{-/-}$ T cells**

| Genotype | Region | Mutation type/ RNApol2 status | Mutation Frequency x10$^{-7}$ | | | Figure |
|---|---|---|---|---|---|---|
| | | | Median | 25% percentile | 75% percentile | |
| *Mlh1*$^{-/-}$ | 3' exons | RNApol2- | 3.73 | 0 | 5.92 | 5E-F |
| | | RNApol2+ | 5.60 | 4.33 | 6.50 | |

| | | | | | | |
|---|---|---|---|---|---|---|
| | 5' exons | RNApol2- | 1.85 | 0 | 2.79 | |
| | | RNApol2+ | 3.83 | 2.29 | 7.40 | |
| | coding | DEL | 2.66 | 1.66 | 3.65 | 4A-C |
| | | INS | 2.70 | 2.08 | 4.55 | |
| | | SNV | 19.2 | 9.64 | 21.4 | |
| | exome | DEL | 7.51 | 6.71 | 7.95 | 2C |
| | | INDEL | 13.7 | 12.7 | 15.2 | 2B |
| | | INS | 6.33 | 5.66 | 7.39 | 2C |
| | | SNV | 16.5 | 11.9 | 23.1 | 2B |
| | noncoding | DEL | 7.23 | 5.23 | 9.05 | 4A-C |
| | | INS | 5.10 | 3.44 | 7.16 | |
| | | SNV | 13.0 | 7.25 | 21.4 | |
| | random | DEL | 2.85 | 2.36 | 3.90 | 3C, E |
| | | INS | 1.36 | 0.467 | 2.20 | |
| | repeat | DEL | 44.2 | 34.0 | 50.8 | |
| | | INS | 58.7 | 51.2 | 70.5 | |
| *Mlh1*$^{+/+}$ | 3' exons | RNApol2- | 1.97 | 0 | 4.36 | 5E-F |
| | | RNApol2+ | 3.39 | 5.74 | 2.41 | |
| | 5' exons | RNApol2- | 1.47 | 2.51 | 4.06 | |
| | | RNApol2+ | 2.27 | 0 | 2.83 | |
| | coding | DEL | 2.60 | 1.22 | 3.47 | 4A-C |
| | | INS | 2.73 | 2.02 | 4.16 | |
| | | SNV | 11.5 | 7.43 | 3.07 | |
| | exome | DEL | 4.09 | 3.61 | 4.60 | 2C |
| | | INDEL | 9.22 | 7.84 | 11.9 | 2B |
| | | INS | 5.05 | 4.62 | 69.2 | 2C |
| | | SNV | 13.1 | 11.1 | 16.6 | 2B |
| | noncoding | DEL | 4.17 | 3.28 | 4.48 | 4A-C |
| | | INS | 4.94 | 3.50 | 5.96 | |
| | | SNV | 10.7 | 6.56 | 16.3 | |
| | random | DEL | 3.33 | 2.71 | 3.79 | 3C, E |
| | | INS | 0.638 | 0.235 | 1.38 | |
| | repeat | DEL | 11.2 | 8.93 | 13.2 | |
| | | INS | 44.7 | 41.1 | 60.0 | |

Mutation frequencies in different genomic regions. Outlier cells (see Methods) are excluded from summary statistics.
